# Cytosine methylation affects the mutability of neighbouring nucleotides in human, Arabidopsis, and rice

**DOI:** 10.1101/764753

**Authors:** Vassili Kusmartsev, Tobias Warnecke

**Affiliations:** Medical Research Council London Institute of Medical Sciences, London, United Kingdom; Institute of Clinical Sciences, Faculty of Medicine, Imperial College London, London, United Kingdom

**Keywords:** Cytosine methylation, mutation, base excision repair, *Arabidopsis thaliana*

## Abstract

Methylated cytosines deaminate at higher rates than unmethylated cytosines and the lesions they produce are repaired less efficiently. As a result, methylated cytosines are mutational hotspots. Here, combining rare polymorphism and base-resolution methylation data in humans, *Arabidopsis thaliana*, and rice (*Oryza sativa*), we present evidence that methylation state affects mutation dynamics not only at the focal cytosine but also at neighbouring nucleotides. In humans, contrary to prior suggestions, we find that nucleotides in the close vicinity (±3nt) of methylated cytosines mutate less frequently. In contrast, methylation is associated with increased neighbourhood mutation risk in *A. thaliana* and rice. The difference in mutation risk associated with methylation is less pronounced further away from the focal CpG, is modulated by regional GC content, and enhanced in heterochromatic regions. Our results are consistent with a model where elevated risk at neighbouring bases is linked to lesion formation at the focal cytosine and subsequent long-patch repair. Our results provide evidence that cytosine methylation has a broader mutational footprints than commonly assumed. They also illustrate that methylation is not intrinsically associated with higher mutation risk for surrounding bases, but that mutagenic effects reflect evolved species-specific and lesion-specific predispositions to elicit error-prone long-patch DNA repair.

## INTRODUCTION

5-methylcytosine (5mC) is found in bacteria, archaea (Blow *et al*. 2016) and diverse eukaryotes, including vertebrates (Goll and Halpern 2011; Li and Zhang 2014), many invertebrates (Regev *et al*. 1998; Bewick *et al*. 2017), plants (Zhang *et al*. 2018), and fungi (Bewick *et al*. 2019). The modification, introduced by methyltransferases that target specific nucleotide contexts, marks the underlying sequence for differential treatment, preventing the binding of some proteins or facilitating recruitment of others, often – though not always – in the context of transcriptional silencing. In mammals, for example, where cytosine methylation is found almost exclusively at CpG dinucleotides, methylation represses transcription at individual promoters and can act in conjunction with histone modifications to establish and maintain larger silent domains, including during X chromosome inactivation (Goto and Monk 1998). In plants, methylation is principally associated with silencing of transposable elements, but can also be found in the bodies of constitutively expressed genes and contributes to dynamic regulation of multiple individual loci (Zhang *et al*. 2018). In both mammals and plants, interfering with methylation is typically deleterious and affects development, survival, or the ability to respond to environmental challenges.

Despite its importance for genome regulation, cytosine methylation comes at a cost: 5mCs are more liable to spontaneous deamination than unmethylated cytosines (Coulondre *et al*. 1978; Duncan and Miller 1980; Wang *et al*. 1982; Ehrlich *et al*. 1986; Zhang and Mathews 1994; Shen *et al*. 1994) and deaminate to thymine rather than uracil. In addition, the resulting T:G mismatches are less efficiently repaired than U:G mismatches (Schmutte *et al*. 1995; Krokan and Bjoras 2013). Methylated cytosines therefore carry the double burden of a higher rate of lesion formation and less efficient repair and, consequently, are more likely to give rise to mutations, specifically C to T transitions (Lutsenko and Bhagwat 1999).

The elevated mutability of 5mC has had and continues to have repercussions for patterns of genetic variation and genome evolution. CpGs are more likely than other dinucleotides to be found polymorphic in mammalian populations (Barker *et al*. 1984; Xia *et al*. 2012) and CpG to TpG changes are disproportionately common amongst variants associated with disease (Cooper and Krawczak 1989; Denissenko *et al*. 1997; Mancini *et al*. 1997; Zemojtel *et al*. 2009; Cooper *et al*. 2010; Zemojtel *et al*. 2011). On the flipside, transitions at CpGs more frequently provide the raw material required for adaptation to novel environments (Stoltzfus and McCandlish 2017; Storz *et al*. 2019). The mutational impact of methylation is also visible over longer evolutionary timescales: CpG to TpG transitions often dominate substitution profiles between species (Ebersberger *et al*. 2002; Hwang and Green 2004) and genomes where CpG methylation is common are depleted of CpGs (Josse *et al*. 1961; Russell *et al*. 1976; Salser 1978; Bird 1980; Simmen 2008). Importantly, higher transition rates at CpGs are also evident in data from parent-child trios (Kong *et al*. 2012; Francioli *et al*. 2015; Rahbari *et al*. 2016; Jónsson *et al*. 2017) somatic mutations in healthy tissues (Hoang *et al*. 2016; Martincorena *et al*. 2018), mutation accumulation lines (Ossowski *et al*. 2010; Lee *et al*. 2012; Weng *et al*. 2019), and when considering rare SNPs (Rahbari *et al*. 2016; Carlson *et al*. 2018), strongly supporting mutational processes as the driving force. Finally, whereas early studies had to rely on CpGs as a (reasonable) proxy for methylation, more recent analyses have tethered elevated rates directly to methylation by integrating base-resolution methylation maps with polymorphism/somatic mutation data and comparing rates of evolution or SNP incidence at methylated and unmethylated CpGs explicitly (Ossowski *et al*. 2010; Mugal and Ellegren 2011; Lee *et al*. 2012; Xia *et al*. 2012; Supek *et al*. 2014; Tomkova *et al*. 2016; Weng *et al*. 2019). There is, in short, overwhelming evidence that cytosine methylation strongly impacts the emergence of novel variants, the spectrum of standing genetic variation, genome-wide base composition, and longer-term patterns of genome evolution.

The focal point in understanding shorter- and longer-term effects of methylation on genome fragility and evolution has, quite naturally, been the methylated cytosine itself. Methylation, however, can cast a longer mutational shadow and affect the rates of lesion formation, recognition and repair beyond the focal cytosine. For specific mutational processes, this has been well documented. Notably, methylation increases UV-induced formation of pyrimidine dimers (Tommasi *et al*. 1997; Ikehata and Ono 2007; Banyasz *et al*. 2016) and slows subsequent repair (Tornaletti and Pfeifer 1996). 5mC also affects the formation and repair of other directly adjacent lesions, including oxidation damage at neighbouring guanines (Tomkova and Schuster-Böckler 2018). But might methylation cast a longer shadow still? Cytosine methylation alters the physico-chemical properties of sequence in which it is embedded, affecting helix stability, rigidity, and dynamics (Szer and Shugar 1966; Collins and Myers 1987; Severin *et al*. 2011; Ngo *et al*. 2016). It can induce slight displacement of the surrounding bases to the minor groove of the helix (Heinemann and Hahn 1992; Marcourt *et al*. 1999; Derreumaux *et al*. 2001), perturb biological processes such as cruciform extrusion (Murchie and Lilley 1989), and might therefore influence damage surveillance and handling more broadly. Further, even if elevated lesion risk is initially limited to the methylated base, neighbouring nucleotides can become collateral damage whenever repair involves excision and re-synthesis around the focal lesion, as is the case for mismatch repair (MMR), nucleotide excision repair (NER) and long-patch modes of base excision repair (BER). The excess mutational risk here can derive, for example, from the use of error-prone polymerases for re-synthesis. Alternatively, it might simply come from the transient generation of single-stranded DNA, which is more sensitive to mutagenic insults or required as a substrate for mutagenic enzymes such as APOBEC deaminases. By monitoring the repair of engineered mismatches in reporter constructs, such excess risk at sites in the vicinity of a focal lesion has been demonstrated, both *in vivo* and *in vitro* (Peña-Diaz *et al*. 2012). But are elevated mutation rates at 5mCs a significant trigger for such events? More generally, can one detect footprints of altered mutability around methylated versus unmethylated cytosines in genomic data?

Previously, Qu and colleagues reported a ∼1.5-fold higher incidence of SNPs ±10bp around methylated compared to unmethylated CpGs in both human and medaka fish (*Oryzias latipes*) (Qu *et al*. 2012), consistent with a role for methylation in increasing the mutability of nucleotides in its vicinity. The observation that, during primate evolution, non-CpG substitution rates positively track the density of CpG dinucleotides in L1 transposons (Walser *et al*. 2008) is further consistent with this scenario. Implicating methylation as the causative force behind increased mutation rate, however, requires careful control of context. Mutation rate varies at multiple scales across the genome, depending on local and regional sequence composition, chromatin state, replication timing, and functional context (Hodgkinson and Eyre-Walker 2011; Ségurel *et al*. 2014; Makova and Hardison 2015). Methylated cytosines are unevenly distributed across these contexts, which might show different mutation rates for reasons unrelated to methylation. For example, in *A. thaliana* and other plants, *de novo* methylation of a sizeable subset of cytosines occurs via a process that specifically targets transposable elements rather than the genome at large (Zhang *et al*. 2018). Taking the non-random distribution of methylated cytosines into account is paramount to dissect whether methylation has left a mark on genome variation and evolution beyond the methylated base itself.

Here, we use data on rare polymorphisms in human, *A. thaliana*, and rice to quantify the mutational effect methylation has on adjacent bases. Rare SNPs constitute a better proxy for mutational processes than common SNPs or substitutions as the latter more strongly reflect longer-term selection and gene conversion (Rahbari *et al*. 2016; Zhu *et al*. 2017). Controlling for sequence context and chromatin state, and considering a range of potential confounders, we find, in contrast to previous results, that methylation is associated with *reduced* SNP incidence at CpG-neighbouring sites in human. In both *A. thaliana* and rice, on the other hand, methylation is positively associated with SNP incidence. In *A. thaliana* and human, excess mutability associated with methylation (or lack of methylation, respectively) appears confined to close neighbours (±3bp) and decays with distance to the methylated site, supporting a mechanism that is contingent on lesion formation at the focal CpG. Our work suggests that methylation casts a longer mutational shadow than commonly assumed, acting in a manner that depends on species-specific coupling between lesion formation and downstream choice of repair pathway.

## RESULTS

### Methylation is associated with decreased mutability of neighbouring bases in humans

To establish whether SNP incidence varies as a function of methylation at nearby CpGs, we combined data from large-scale surveys of population genomic variation with base-resolution methylation data. For human, we defined methylated (unmethylated) sites as those with >70% (<20%) methylated reads in H1 human embryonic stem cells (hESC) (Lister *et al*. 2009), previously shown to be a reasonable proxy for germline methylation (Prendergast *et al*. 2014; Supek *et al*. 2014). Analysis was limited to sites covered by at least ten reads (see Methods for further details). As in prior work (Supek *et al*. 2014), we then paired methylated and unmethylated CpGs according to multiple criteria, which were applied simultaneously:

First, as mutation rate strongly varies with local sequence context (Blake *et al*. 1992; Zhao and Boerwinkle 2002; Hwang and Green 2004; Carlson *et al*. 2018), we required the four nucleotides either side (±4bp) of the focal CpG to be the same. Matching the sequence context in this manner also controls for local GC content, previously shown to correlate inversely with CpG mutability (as further discussed below), and sequence complexity, which is an important determinant of indel formation propensity. As heptanucleotide context was previously shown to account for more than 80% of variability in mutation rates in humans (Aggarwala and Voight 2016), we did not extend matching to even longer sequence motifs, which would have drastically reduced sample size.

Second, we required each member of the methylated/unmethylated pair to be in the same chromatin state as defined by a widely used hidden Markov model for H1, which integrates signals from multiple histone marks, methylation and DNA accessibility (Ho *et al*. 2014) (see Methods). Matching by chromatin state is important because mutation rates vary substantially with chromatin environment (Schuster-Böckler and Lehner 2012; Makova and Hardison 2015). In particular, heterochromatic regions accumulate more mutations compared to euchromatic, actively transcribed regions (Schuster-Böckler and Lehner 2012), which are generally more accessible to or specifically targeted by DNA repair machinery (Supek and Ben Lehner 2015; Frigola *et al*. 2017). Chromatin states also capture other determinants of mutation rate heterogeneity, including replication timing and transcriptional activity, the latter being important in the context of this work because deamination risk is more than two orders of magnitude higher in single-compared to double-stranded DNA (Frederico *et al*. 1990).

Requiring the same nucleotide and chromatin context, choosing the closest available match along the same chromosome, and excluding sequence contexts with CpGs other than the focal CpG, we obtained 60,589 pairs of matched sites. At each base surrounding the focal CpG, we then calculated the incidence of singleton SNPs as observed across 15,708 whole genomes from the gnomAD database (Lek *et al*. 2016; Karczewski *et al*. 2019) (see Methods). To avoid looking at compound effects of clustered methylation sites and to allow comparison with previous studies (see below), any SNP that was a T to C transition at a TpG or an A to G transition at a CpA, was excluded. Applying this protocol, we find a reduced incidence of SNPs adjacent to methylated CpGs compared to unmethylated CpGs, illustrated as the relative mutational risk associated with methylation, *RR_met_*, in Figure 1A. Across all sites (±1bp to ±3bp from the focal CpG) and mutation types (transitions and transversion), *RR_met_* is 0.886 (P=2.73*10^−21^, Z-test for proportions). In other words, there are 886 SNPs in the ±3bp neighbourhood of methylated CpGs for every 1000 SNPs surrounding unmethylated CpGs.

**Figure 1.**
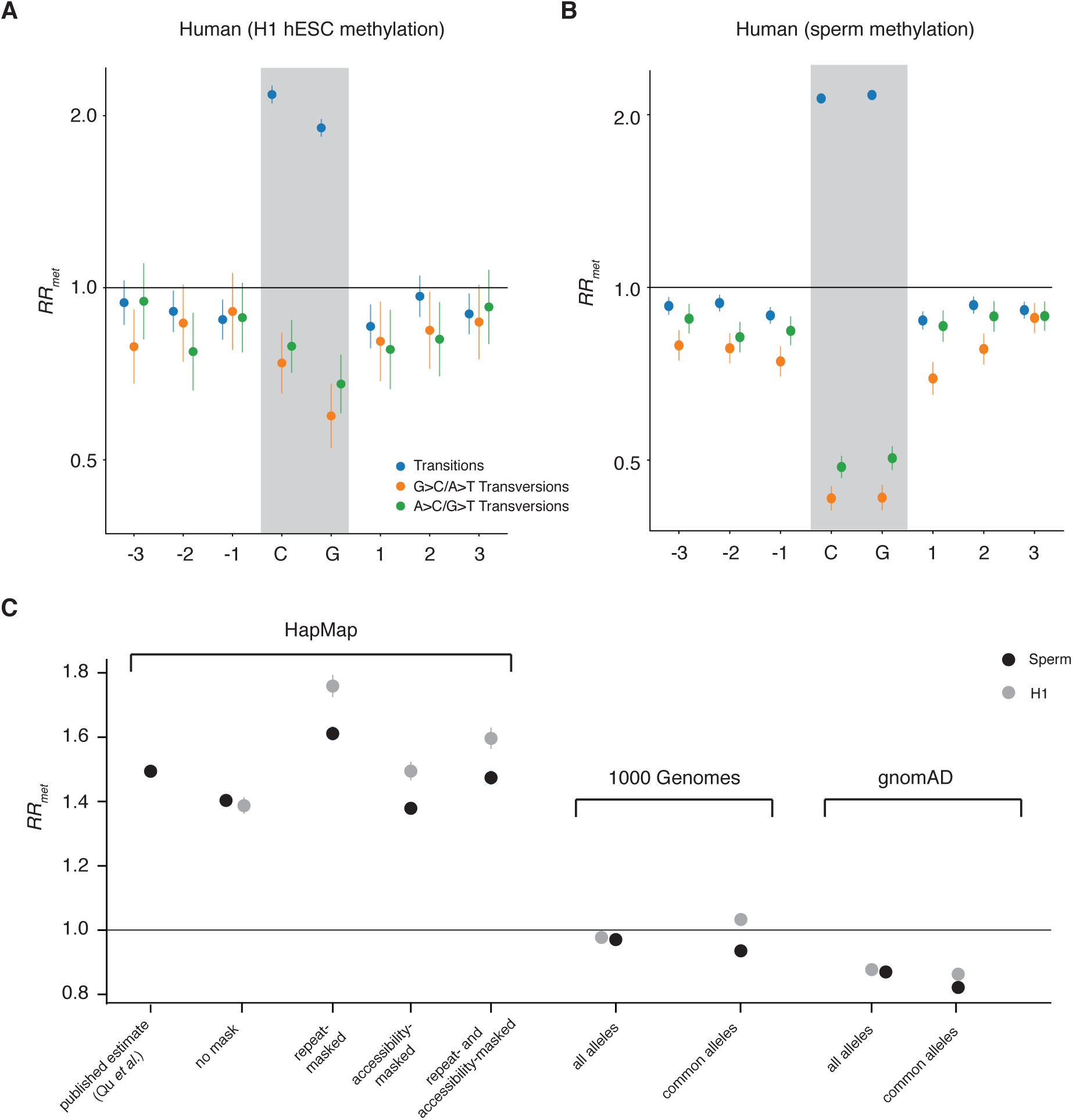
The relative mutational risk of methylation (*RR_met_*) at CpGs and neighbouring nucleotides in human, when site pairing is based on methylation data from **(A)** H1 hESCs or **(B)** sperm and considering singleton SNPs from the gnomAD database. **(C)** The impact of analytical and dataset choice on *RR_met_*estimates. Polymorphism data from the HapMap project (CEU) consistently lead to *RR_met_*estimates >1, which are robust to inclusion of SNPs from poorly accessible or repeat regions. *RR_met_*estimates are below one when considering SNPs from the 1000 Genomes Project or gnomAD. Common alleles are defined as alleles with a frequency of ≥5% in the population sample. All P<10-8. Vertical bars are confidence intervals on *RR_met_*computed using the delta method.

### Prior contradictory results are linked to the use of HapMap variation data

The above results appear to contradict a prior analysis of genome-wide human polymorphism and methylation data, which found more mutations in the vicinity of methylated sites (Qu *et al*. 2012). The analytical pipeline of Qu and colleagues (hereafter simply referred to as Qu) exhibits several potentially important differences to our approach. First, Qu used methylation data from sperm (Molaro *et al*. 2012) rather than H1 hESC (Lister *et al*. 2009). Second, they considered SNPs of any frequency from the HapMap project (CEU population) while we consider rare SNPs from the much larger gnomAD database. Third, whereas we pursue the matching approach described above, Qu assigned nucleotides in the vicinity (±10bp) of a given CpG to one of two categories: joining blocks of overlapping CpG dinucleotides, they averaged methylation levels across CpGs in the resulting larger block and considered methylated (unmethylated) blocks to be those with overall CpG methylation levels ≥80% (≤20%). They then computed an overall SNP incidence rate for each category. Finally, Qu analysed sites with ≥5 reads.

To understand which – if any – of these analytical choices explain the discrepancy, we started by reimplemented our matching approach with the sperm methylation data used by Qu. We obtained very similar results (*RR_met_*=0.877, P=1.68*10^−160^, Figure 1B), suggesting that the different methylation datasets are not the source of the discrepancy. We also obtained very similar results when considering sites with ≥5 instead of ≥10 reads (overall RR_met_=0.889, P=3.19*10^−81^), which increases samples size to 162,156 pairs. Unsurprisingly, given that the distribution of methylation stoichiometries exhibits two modes at the extremes (i.e. towards 0% and 100% methylated, see Supek *et al*. 2014), there is also no substantive change when we require ≥80% rather than ≥70% read support to call a site methylated (not shown). Next, we sought to reproduce the original finding by implementing the approach of Qu in full, using HapMap (CEU) polymorphisms, sperm methylation data, sites covered by ≥5 read, a threshold of ≥80% for calling methylated sites, and calculating *RR_met_* as the SNP incidence in blocks around focal CpGs as described above. Doing so, we can replicate their results, at least qualitatively (*RR_met_*=1.4, Figure 1C). Using H1 instead of sperm data again yielded very similar results (Figure 1C), re-confirming that differences in methylation data are not pertinent. We then checked whether inclusion of repeats or regions associated with less reliable SNP calls, might explain the difference. Excluding repeats or poorly accessible regions, as defined by the 1000 Genomes project (see Methods), however, has limited effect, with *RR_met_* consistently >1 (Figure 1C). In contrast, we obtain dramatically different results when substituting HapMap (CEU) polymorphisms for singleton SNPs from the 1000 Genomes project (overall *RR_met_*=0.971) or gnomAD (overall *RR_met_*=0.87, Figure 1C).

One key difference between the variation datasets above is the relative prominence of common alleles, which is much greater in HapMap. One might therefore reasonably hypothesize that different *RR_met_* estimates in the HapMap, 1000 Genomes, and gnomAD data are owing to differences in average allele frequency, perhaps because common alleles are a poorer proxy for mutational processes, reflecting selection and gene conversion to a greater extent than rare SNPs. However, allele frequency appears to be only a minor factor. We obtain similar results (i.e. *RR_met_*<1) when we threshold 1000 Genomes and gnomAD data to only include SNPs present at ≥5% frequency in the given sample. We therefore conclude that difference in SNP quality/mapping in the original HapMap data likely underlie difference between our results and those of Qu. As rare SNPs provide better proxy for mutational processes, we further conclude that there is no evidence that methylation is associated with increased mutability at adjacent sites. Rather, cytosine methylation is associated with a significant reduction in the incidence of mutations in the neighbourhood of CpGs in humans. As we discuss in greater depth below, this is consistent with recent experimental data.

### Methylation is associated with increased SNP incidence in plants

In contrast to humans, we find SNP-approximated mutability of CpG-neighbouring bases to be positively associated with methylation in two plants: *A. thaliana* and rice. Applying the same matching protocol (and confining analysis to repeat-masked sequence, see Methods), we find an overall *RR_met_* of 1.28 (P=2.06*10^−89^) in *A. thaliana* and 1.31 in *O. sativa* (P=5.43*10^−15^), where estimates are noisier due to relatively smaller number of matched pairs (Figure 2A; N=121,774 pairs in *A. thaliana*, N=42,779 pairs in *O. sativa*). The strongest increase in mutational risk was associated with C to T transition SNPs (*RR_met_*=1.38 in *A. thaliana*; *RR_met_*=1.71 in *O. sativa*). Interestingly, *RR_met_* appears to level off as a function of distance from the focal CpG, with the greatest deviation from random expectation (*RR_met_*=1) at the nucleotide directly adjacent to the CpG. A similar (albeit inverted) trend is also evident in humans (Figure 2A, Figure 1A/B).

**Figure 2.**
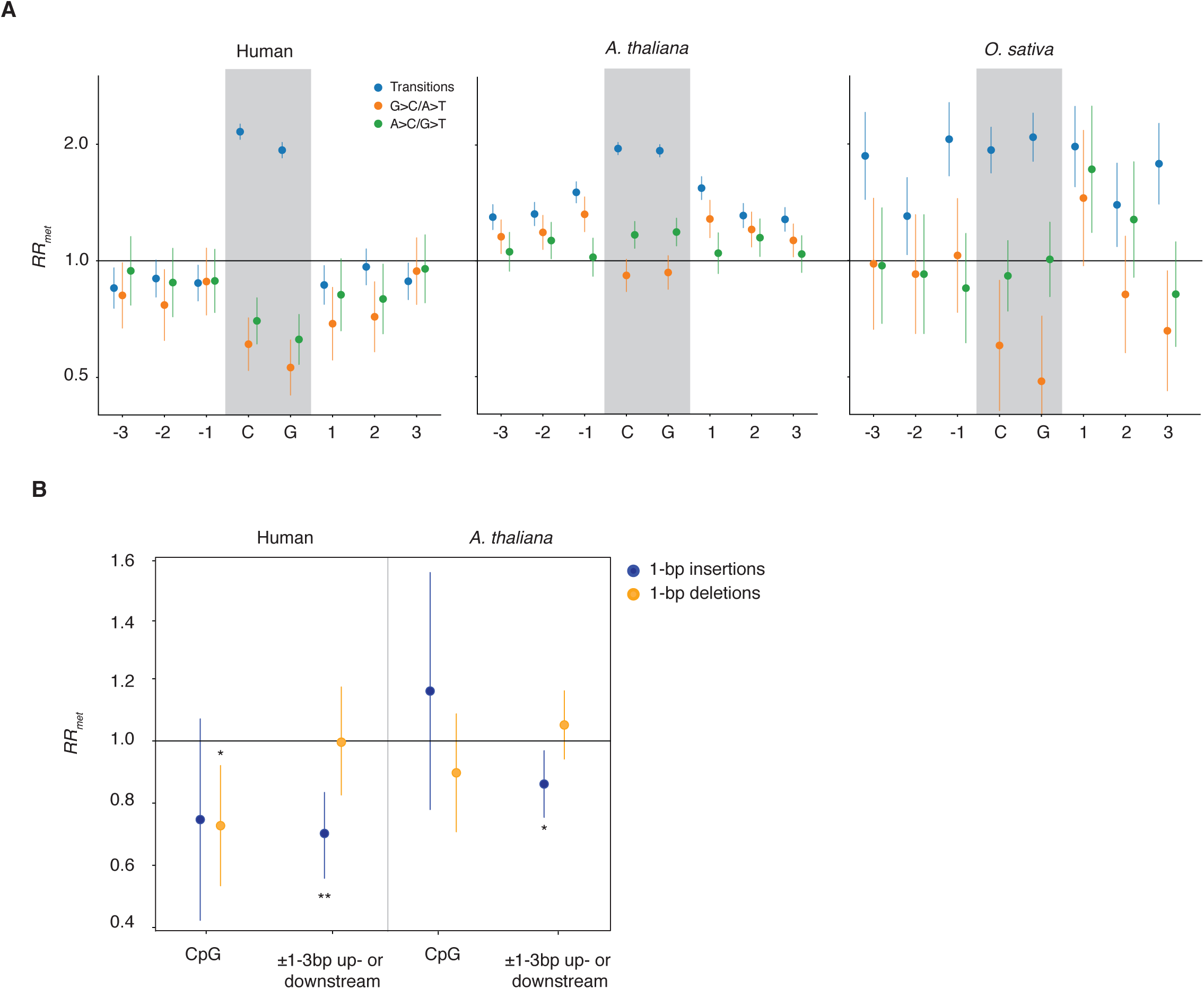
(**A**) The relative mutational risk of methylation (*RR_met_*) at CpGs and neighbouring nucleotides in human, *A. thaliana*, and rice. For human, pairing is based on H1 hESC methylation data. In contrast to Figure 1, only SNPs in repeat-masked sequence was included for all species. (**B**) *RR_met_* as applied to indels in human and *A. thaliana*. Vertical bars are confidence intervals on *RR_met_* computed using the delta method. *P<0.05, **P<0.005

### Methylation is associated with an altered incidence of insertions and deletions

With a view to understanding the mechanism of altered neighbourhood mutability, we also investigated whether the presence of methylation affected the rates of insertions and deletions (indels) around the focal CpG, using the same matched pairs as above. As for SNPs, we focus on singleton indels and also confine analysis to single-base insertions and deletions, thus ensuring that the sequence context of the indel is comparable. As indels are rarer than SNPs, we compute only two *RR_met_* estimates, for the CpG itself and all neighbouring bases upstream (±1-3bp) combined. In humans, echoing results from SNPs, we find a reduced incidence of indels for methylated sites, both at the CpG itself and the neighbouring sequence (Figure 2B). In *A. thaliana, RR_met_* is higher across the board, with a tendency for *RR_met_*>1 for deletions in CpG-proximal sites. There were too few events to carry out an informative analysis of indel patterns in rice.

### Increased mutability around methylated sites in plants is not limited to the CpG context

As methylation in plants is not limited to CpGs but also occurs in CpHpG and CpHpH contexts (where H=A, C, or T), we independently collated matched pairs for these contexts. In both cases, there is a tendency for *RR_met_* to be greater than 1 (Figure S1). However, as methylation at CpHpG and CpHpH sites is rarer (6% and 1.5% compared to 24% for CpG in *A. thaliana*, 21% and 2.2% compared to 59% for CpG in *O. sativa*) and more strongly skewed to certain functional contexts, sample sizes are much smaller. We therefore focus on CpGs below, but note that elevated *RR_met_* across all contexts suggest that differential risk is likely not associated with machinery that acts exclusively at CpGs versus other contexts.

### Methylation-associated changes in mutability are not sequence- or context-specific

To identify potential drivers of methylation-associated differences in mutability (and pinpoint difference between plants and humans), we stratified *RR_met_* by sequence context, local/regional GC content, and chromatin state. First, we note that higher mutability around methylated CpGs is observed across most central sub-contexts (CpG ±1bp) that are sufficiently common (present in more than 20 pairs) to allow independent assessment of SNP rates (Figure 3A). This suggests that, although there is variability associated with the surrounding sequence, the effect is not driven by a specific nucleotide context or set of contexts. This also provides a first pointer that differential context composition does not explain why observations in plants differ from those in human. To rule this out explicitly, we subsampled matched pairs to achieve an identical representation of nucleotide contexts in human and *A. thaliana*. Doing so, *RR_met_* remains >1 in *A. thaliana* and <1 in human (Figure S2).

**Figure 3.**
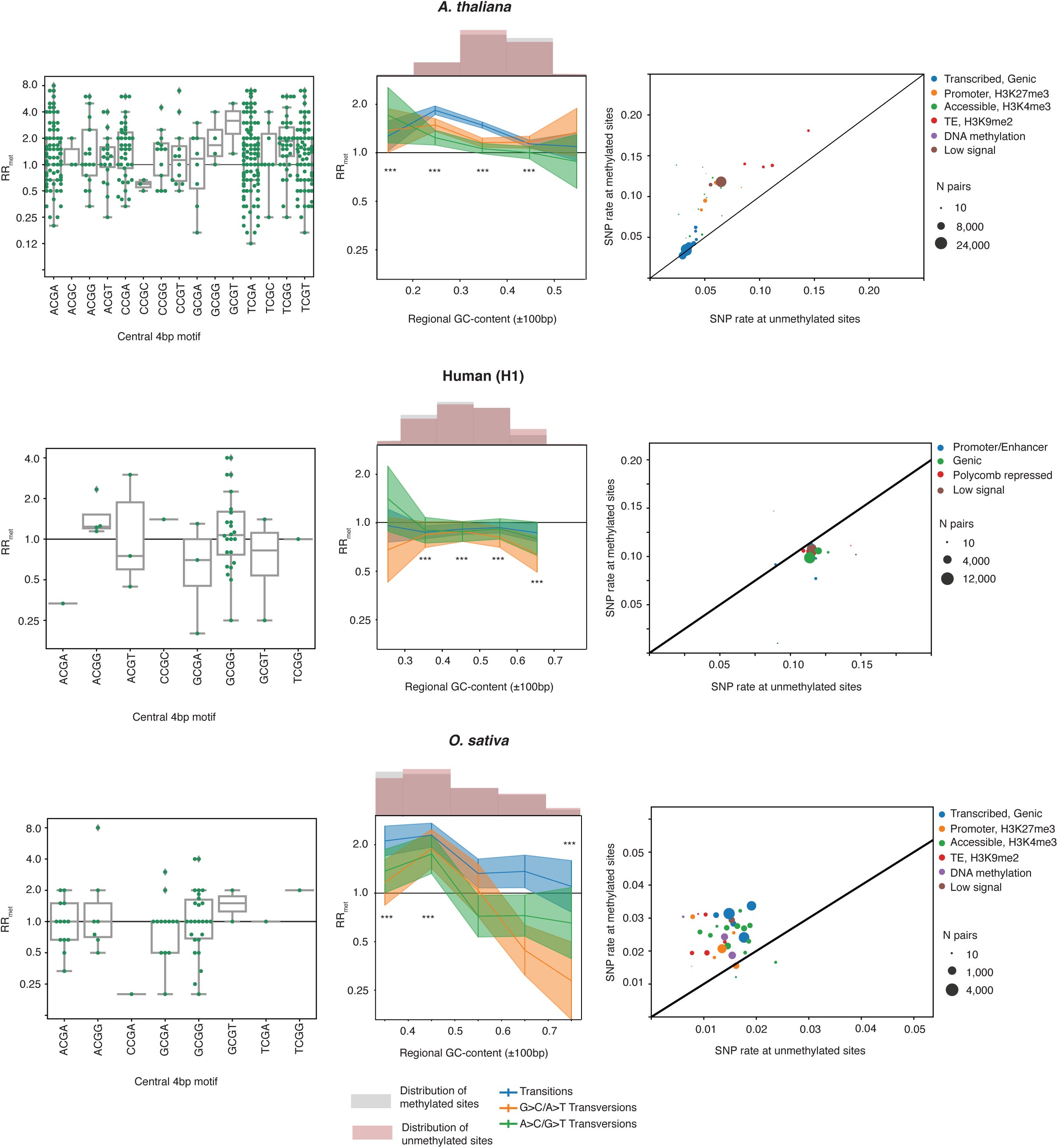
The relative mutational risk of methylation (*RR_met_*) at CpG-neighbouring nucleotides in human, *A. thaliana*, and rice as a function of internal (±1bp) sequence context (left-hand panels), regional (±100bp) GC content (central panels), and chromatin state (right-hand panels). To calculate *RR_met_* for motifs with a given internal sequence context, only motifs with a minimum number of 20 pairs were considered. To calculate dependency on GC content, methylated and unmethylated members of a given pair were independently binned based on their regional GC content and *RR_met_* calculated based on these bins. The distribution of regional GC content for methylated and unmethylated sites is almost identical. ***P<0.001. Vertical bars are confidence intervals on *RR_met_* computed using the delta method. *RR_met_* for different chromatin states, deconstructed into SNP rates at methylated and unmethylated sites across pairs, is colour-coded according to broader overarching categories of similar states whose principal features are highlighted. See Figure S4 for fully annotated states.

We then considered to what extent *RR_met_* is influenced by or robust to local and regional GC content. GC content has previously been suggested to impact focal deamination rates as DNA duplexes of high GC content are less likely to spontaneously become single-stranded (Leroy *et al*. 1988). In line with this model, observed CpG content was previously found to be lower than expected in regions of low GC but not in regions of high GC content (Adams and Eason 1984) and substitution rates at CpGs in primates and Arabidopsis correlate inversely with GC content (Fryxell 2004; Leah J DeRose-Wilson 2007). Considering local (motif-wide) and regional (±100bp either side of the focal CpG) GC content, we find that *RR_met_* in plants is highest at low-to-intermediate GC content and becomes less pronounced at higher GC content (Figure 3B, Figure S3). This is consistent with a collateral damage model of mutagenesis, where initial deamination events at methylated focal CpGs put neighbouring sites at risk and do so more frequently in a low GC context, where spontaneous deamination is more likely to occur. In humans, we observe no such relationship, in line with the absence of 5mC deamination as the driver of mutational liability.

Next, we considered variability in *RR_met_* across different chromatin states. Again, we find consistent signal *RR_met_* >1 in plants (Figure 3C, Figure S4). However, some chromatin states emit much stronger signals than others. Notably, chromatin states characterized by the presence of transposable elements and/or histone marks that are generally associated with silencing (H3K27me3, H3K9me2), not only have higher absolute SNP incidence but also notably higher *RR_met_* compared to transcribed genic sequence. Some chromatin states marked by high accessibility (DNase hypersensitivity) also have high *RR_met_*. However, this is only true for states (e.g. promoters) outside of transcribed regions, where the methylation-associated risk of mutation appears to be strongly reduced, perhaps as a result of transcription-coupled repair.

### Sites accessible to the (de)methylation machinery show reduced relative risk

In both mammals and plants, methylated cytosines are subject to active demethylation. In mammals, ten-eleven translocation (TET) enzymes oxidize 5mC residues to yield 5-hydroxymethylcytosine, which – along with further-oxidized derivatives – can be excised by DNA glycosylases like TDG. In plants, active demethylation is more direct. Different glycosylases – such as DEMETER (DME) and REPRESSOR OF SILENCING 1 (ROS1) in *A. thaliana* – target and excise 5mC itself (Zhu 2009). Both direct or indirect demethylation pathways, being reliant on DNA glycosylase activity to generate abasic sites, effectively constitute instances of programmed lesion formation. For mammals, our lab and others have previously shown that cytosines that spend a larger fraction of their time in a hydroxymethylated state are subject to different mutation dynamics (Supek *et al*. 2014; Tomkova *et al*. 2016). In a similar vein, we wanted to know whether repeated activity of the *A. thaliana* demethylation machinery might similarly be associated with altered mutagenesis, which might also affect neighbouring sites. In particular, we hypothesized that sites that undergo frequent methylation/demethylation/remethylation cycles might suffer from a greater cumulative risk of mutation if repair following glycosylase activity – thought to be principally BER – is mutagenic. Sites that undergo such cycles are not uncommon in *A. thaliana*. ROS1 in particular has been implicated in preventing, through ongoing active removal of 5mCs, the spread of methylation from transposable elements into active genes (Zhang *et al*. 2018).

We therefore considered *RR_met_* in the context ROS1 activity, classifying sites as ROS1 targets if they showed increased methylation in a *ros1* knock-out strain (see Methods). Similarly, we considered sites to be regular targets of the RNA-directed DNA methylation (RdDM) machinery, which is responsible for *de novo* methylation, if they showed decreased methylation in strains deleted for *nrpd1,* a polymerase IV subunit critical for methylation.

Here, in order to include effects of ROS1-targeted sites, we considered changes in methylation in *nrpd1/ros1* double mutants compared to the *ros1* mutant. We paired sites by sequence context and by ROS1/NRPD1 target status. As *RR_met_* is >1 throughout different chromatin contexts, we jettison chromatin state pairing to maintain a sufficient samples size for analysis. We find that sites targeted by ROS1 experience a substantially lower excess mutation risk associated with methylation than sites that do not experience increased methylation upon *ros1* knock-out (Figure 4A). We observe very similar results for *nrpd1* (Figure 4A). Post hoc analyses suggests that these differences are also observed when chromatin state is controlled for (not shown). We note that this effect is quantitative: sites that experience greater gain (loss) in methylation in *ros1* (*nrpd1*) knock-outs, show lower *RR_met_* (Figure 4B). Finally, cytosines that are dependent on the chromatin remodeler DDM1 for methylation show higher *RR_met_* than sites where methylation can be established and subsequently maintained solely through RdDM and maintenance methyltransferase activity (Figure 4A, see Methods for how DDM1 dependency was defined).

**Figure 4.**
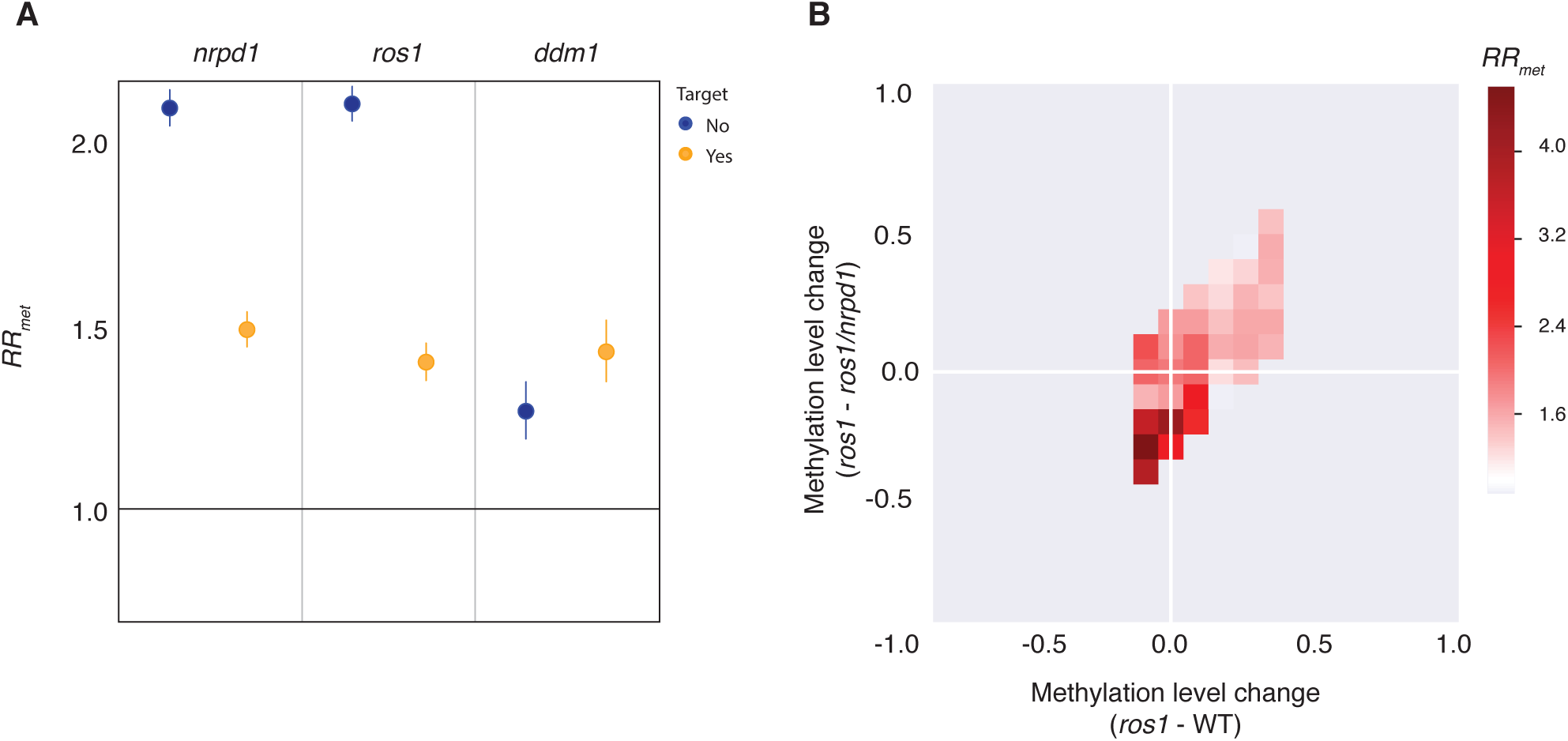
(**A**) The relative mutational risk of methylation (*RR_met_*) at CpGs and neighbouring nucleotides in *A. thaliana* as a function of whether a given site is targeted by RNA-directed DNA methylation (which involves *nrpd1*), the DNA glycosylase ROS1, or the chromatin remodeler DDM1 (see main text for how targets and non-targets were defined). All P<10^−60^. Vertical bars are confidence intervals on *RR_met_* computed using the delta method. (**B**) *RR_met_* as a function of relative methylation change in *ros1* deletion mutants compared to the corresponding wildtype strain and compared to the *nrpd1/ros1* double mutants. Sites with more methylation in *ros1* compared to the WT are sites of high ROS1 activity, where methylation is normally erased by ROS1 but increases in the *ros1* knock-out. Sites with more methylation in *ros1* compared to *nrpd1/ros1* are sites where the RNA-directed DNA methylation machinery is active, i.e. where deletion of *nrpd1* leads to further loss of methylation.

Taken together, these results are inconsistent with a model where higher mutability of neighbouring bases is a consequence of frequent methylation/demethylation cycles. Rather, they reinforce the impression that *RR_met_* is enhanced in regions that are less accessible to machinery that affects methylation, demethylation and presumably demethylation-coupled repair.

## DISCUSSION

Our results suggest that methylation state affects mutability of neighbouring nucleotides in human, *A. thaliana* and rice. Notably, we find *RR_met_*>1 in plants but *RR_met_*<1 in human, which is not explained by differential representation of specific sequence or chromatin contexts. This suggests that differential effects of methylation across species cannot be traced back to difference between methylated/unmethylated sequence with respect to their biophysics, which is species-invariant, but must instead be caused by differential responses by cellular machinery to methylation or methylation-associated lesions. We therefore suggest that *RR_met_* is not linked to methylation per se, but reflects the probability of lesion formation, the nature of the resulting lesion, and how the cellular machinery deals with that lesion.

In human and *A. thaliana*, the spatial pattern of *RR_met_* around the focal CpG returns relatively quickly towards the baseline. Even though our matching approach curtails examination of longer-range effects, this suggests that the mutational effects are relatively local. We think that this locally confined signal is suggestive of BER, which has previously been tipped as a potential culprit behind neighbourhood mutation effects (Qu *et al*. 2012). Experiments in cell extracts have demonstrated the existence of different flavours of BER in both humans and plants (Córdoba-Cañero *et al*. 2009; Martínez-Macías *et al*. 2013), which differ in the number of bases excised and re-synthesized during the repair process. In short-patch BER (SP-BER) only a single nucleotide is added whereas during long-patch BER (LP-BER) multiple bases are excised and re-synthesized. In *A. thaliana*, tracking repair at U:G mismatches, repair tract lengths of up to 3bp have been observed (Córdoba-Cañero *et al*. 2009; Martínez-Macías *et al*. 2013). In mammals, tracts removed during LP-BER are similarly short, 2-13bp (Fortini and Dogliotti 2007; Krokan and Bjoras 2013). This contrasts with nucleotide excision and mismatch repair, where excision tract lengths are typically much longer, from >20bp for NER (Sancar 1996) to several hundred bases or more during MMR (Fang and Modrich 1993). The involvement of LP-BER is also consistent with increased single-nucleotide deletion rates (Bennett *et al*. 2001; Lyons and O’Brien 2010).

But why would trends be inverted for human and plants? We posit that *RR_met_* reflects the combination of two distinct risk factors: the pathway chosen downstream of a given lesion (which determines sign; e.g. whether U:G or T:G is handled more efficiently), and the relative risk of lesion formation (which should be higher for T:G and affects amplitude). In humans, while some lesions, including 8-oxoguanine, are thought to mostly trigger SP-BER (Fortini *et al*. 1999), others have been associated with LP-BER. Importantly, this includes U:G mismatches, that result from deamination of unmethylated cytosines. Studying mouse embryonic fibroblasts, Bennett *et al*. showed that, where uracil removal is triggered by uracil DNA glycosylases (UNG), subsequent re-synthesis exceeded a single nucleotide in 80% of cases, suggesting that UNG, perhaps by virtue of interacting with PCNA, biases downstream repair pathway choice towards long-patch repair (Bennett *et al*. 2001; Fortini and Dogliotti 2007).

Importantly, such “uracil-initiated” (Bennett *et al*. 2001) LP-BER has already been associated with higher mutation risk at neighbouring sites: Chen and colleagues introduced mismatches into an SV40 episome capable of replicating in human cells to monitor mutagenic effects either side of the mismatch. While they found that both T:G and U:G mismatches were associated with collateral damage to the neighbourhood, liability was ∼7-fold higher for U:G (Chen *et al*. 2014). Thus, even though, in a physiological context, the rate of lesion formation might be higher for T:G, the mutagenesis risk associated with repair might be greater for U:G. This is in line with our findings. The precise mechanism(s) behind increased rates surrounding unmethylated CpGs in our data, however, remains unclear. Chen *et al*. showed that both BER and MMR were required for elevated mutability and proposed a model inspired by events during somatic hypermutation at immunoglobulin genes, where BER-generated lesions are hijacked by the MMR machinery, as previously demonstrated for U:G repair (Schanz *et al*. 2009; Peña-Diaz *et al*. 2012) and known to occur in the context of active demethylation (Grin and Ishchenko 2016). Specifically, they suggested that excess mutability in their experimental model is consistent with APOBEC enzymes tagging along with the MMR machinery to attack and deaminate single-stranded cytosines. In our data, however, we find no enrichment of the tell-tale APOBEC mutational signature (C to U changes in a TpCpN context). APOBEC-related excess mutational risk might therefore be a specific manifestation of a more general liability of being more fragile in a single-stranded state. In plants, both LP- and SP- BER have been observed in *A. thaliana*. How different lesions (or associated glycosylases) predispose to either short- or long-patch repair remains poorly understood (Lee *et al*. 2014), but – as the repertoire of glycosylases differs substantially between humans and plants – it is certainly conceivable that U:G and T:G mismatches might be associated with a different propensity for LP-versus SP-BER to what is seen in humans.

We speculate in this regard that choice of repair (sub-)pathways in different species – and the ensuing mutational burden – might be evolved rather than random. Methylation in plants is more tightly linked to transposable elements than it is in mammals. Consequently, there is likely less selection in plants to prevent lesions at and around methylated sites. Where methylated CpGs are located in functionally important regions (gene bodies), they are relatively less affected, perhaps because of transcription-coupled repair (Figure 3C). In mammals, on the other hand, there are more cases where methylation-based silencing is transient and the underlying sequence remains important, notably in the context of X inactivation. Elevated mutation loads in the neighbourhood of methylated cytosines might therefore be much less well tolerated in mammals, so that better repair of these lesions evolved. It is interesting to note in this regard that, during post-replicative repair in mammals, G:T mismatches in the context of hemi-methylated sites are preferentially corrected to G:C with high (∼90%) efficiency (Brown and Jiricny 1987; Bill *et al*. 1998), perhaps reflective of an evolved bias to counter frequent deamination at 5mC. In plants, no such bias has been observed although direct evidence is limited to tobacco protoblasts (Inamdar *et al*. 1992) and this issue deserves further investigation.

Irrespective of the mechanistic underpinnings of altered methylation-associated mutability, which remain to be resolved, our study provides strong evidence that the mutational impact of methylation extends beyond the methylated cytosine itself and shapes the emergence of novel variants across the genomes of different eukaryotes.

## METHODS

### Analysis of relative mutational risk associated with methylation in human

To isolate the specific impact of methylation on SNP incidence, we pursued a matched pairs approach, broadly as previously described (Supek *et al*. 2014). First, CpGs in the human genome were classified as methylated or unmethylated based on base-resolution methylation information in H1 human embryonic stem cells (Lister *et al*. 2009) (http://neomorph.salk.edu/human_methylome/data.html). Methylated sites were defined as cytosines with a ≥0.7 ratio of methylated to unmethylated reads; unmethylated sites as those with a ratio ≤0.2. CpGs with intermediate methylation (0.2>x<0.7) levels were excluded from further analysis. Note that, as methylation stoichiometry is strongly bimodal, this excludes relatively few sites (see Supek *et al*. 2014 for details). To provide robust classification of methylated/unmethylated sites, we only considered CpGs with ≥10 read coverage in the bisulfite sequencing data. As the X chromosome is differentially affected by methylation in males and females and because mutations that arise on sex chromosomes are subject to distinct evolutionary regimes, analysis was confined to autosomes. To enable intersection with polymorphism data, coordinates for all eligible methylated/unmethylated CpG sites were converted to hg19 using the UCSC LiftOver utility (https://genome.ucsc.edu/cgi-bin/hgLiftOver). Each unmethylated CpG was then paired with the nearest methylated CpG that matched the following criteria: a) identical ±4bp flanking sequence context in the reconstructed ancestral human genome (ftp://ftp.1000genomes.ebi.ac.uk/vol1/ftp/phase1/analysis_results/supporting/ancestral_alignments/), b) identical chromatin state as defined by a hidden Markov model for H1 (https://personal.broadinstitute.org/anshul/projects/wfh/ihmm/chmmBED/human_H1/), c) located on the same chromosome. Subsequently, any pairs whose flanking sequence context contained one or more additional CpG dinucleotides were removed to avoid possible confounding effects of proximal CpGs on local mutation risk. Further, pairs that included CpGs not captured by the H1 HMM were also removed. For the surviving pairs of methylated and unmethylated CpGs, we then calculated the incidence of singleton SNPs from the gnomAD database (Karczewski *et al*. 2019) (https://gnomad.broadinstitute.org/). In calculating the incidence of transition SNPs, we removed T to C SNPs that had occurred in the TpG context and A to G SNPs that had occurred in the CpA context, principally to enable comparison to the analysis of Qu et al, where – given the data available at the time – it was prudent to exclude these sites because of potential polarization errors. Note that, above, we only consider SNPs at positions ±3bp from the focal CpG despite pairing for ±4bp. This is because at the ±4 position, the ±5 base is unknown, and so including this position could lead to skews due to unknown neighbouring bases.

### Analysis of relative mutational risk associated with methylation in Arabidopsis and rice

The protocol outlined above was also applied to *A. thaliana* and rice. Classification of CpGs (but also CHG and CHH sites) into methylated and unmethylated sites was based on bisulfite data of Ws0 global stage seed for *A. thaliana* (GSM1664380) (Lin *et al*. 2017), and data from 3-week old leaf tissue in rice (GSM1039487) (Stroud *et al*. 2013). As highlighted above, to calculate rare SNP incidence, we considered homozygous SNPs confined to a single accession in the selfing *A. thaliana* as singletons. HMM-defined chromatin states for both *A. thaliana* and rice were taken from the Plant Chromatin State Database (Liu *et al*. 2018) (http://systemsbiology.cau.edu.cn/chromstates/). The 1001 Arabidopsis genomes (https://1001genomes.org/data/GMI-MPI/releases/v3.1/) and the 3000 rice genomes projects (http://snp-seek.irri.org/), respectively, were the sources of polymorphism data. Repeat sequences in both plants, as well as human, were masked using RepeatMasker v4.0.8 (www.repeatmasker.org).

### Methylation/demethylation mutants

We obtained base-resolution methylation data for knock-out mutants of *ros1* (GSM1859475), *nrpd1* (GSM1859476), and a *ros1*/*nrpd1* (GSM1859478) double knock-out (Wierzbicki *et al*. 2012) as well as for DDM (GSM1014117) and DRD (GSM1014120) mutants (Zemach *et al*. 2013). Base calls were lifted over to TAIR10 as required. Sites affected by a given deletion were classified as follows: a site targeted by ROS1 was one where methylation was greater in the *ros1* mutant than in the WT; a site targeted by NRPD1 was one where methylation was lower in the *nrpd1/ros1* double mutant compared to the *ros1* deletion strain; a site requiring DDM for methylation was one where the *ddm* mutant had lower methylation and the *drd* mutant had no effect.

### Replicating prior estimates of relative mutational risk associated with methylation

To track down divergent *RR_met_* estimates compared to the study of Qu, we additionally considered base-resolution bisulfite sequencing data from sperm (Molaro *et al*. 2012) (GSE30340), lifted over to hg19, and re-implemented their original protocol based on the published methods and feedback provided by the lead author, Wei Qu. This included consideration of sites with ≥5 read coverage, with unmethylated/methylated CpGs defined as those with methylation stoichiometries of ≥0.8 and ≤0.2, respectively. Nucleotides flanking the focal CpG (±10bp) were considered part of methylated or unmethylated “blocks” based on the methylation status of the focal CpG. These site blocks, defined according to either H1 or sperm methylation data, were then overlapped with polymorphism data, either hg19-converted HapMap (ftp://ftp.1000genomes.ebi.ac.uk/vol1/ftp/release/20130502/), 1000 Genomes (ftp://ftp.1000genomes.ebi.ac.uk/vol1/ftp/release/20130502/), or gnomAD (https://gnomad.broadinstitute.org/) (Karczewski *et al*. 2019). As in the original study, SNPs that occurred in CpG/TpG/CpA dinucleotides were excluded.

## Acknowledgements

The authors are grateful to Wei Qu for help in replicating her original protocol and members of the Molecular Systems group for discussions. This work was supported by UK Medical Research Council core funding to TW.

## Author contributions

VK carried out all analyses. TW conceived the study. VK and TW designed analyses, interpreted results, and wrote the manuscript.

---

**Figure S1.**
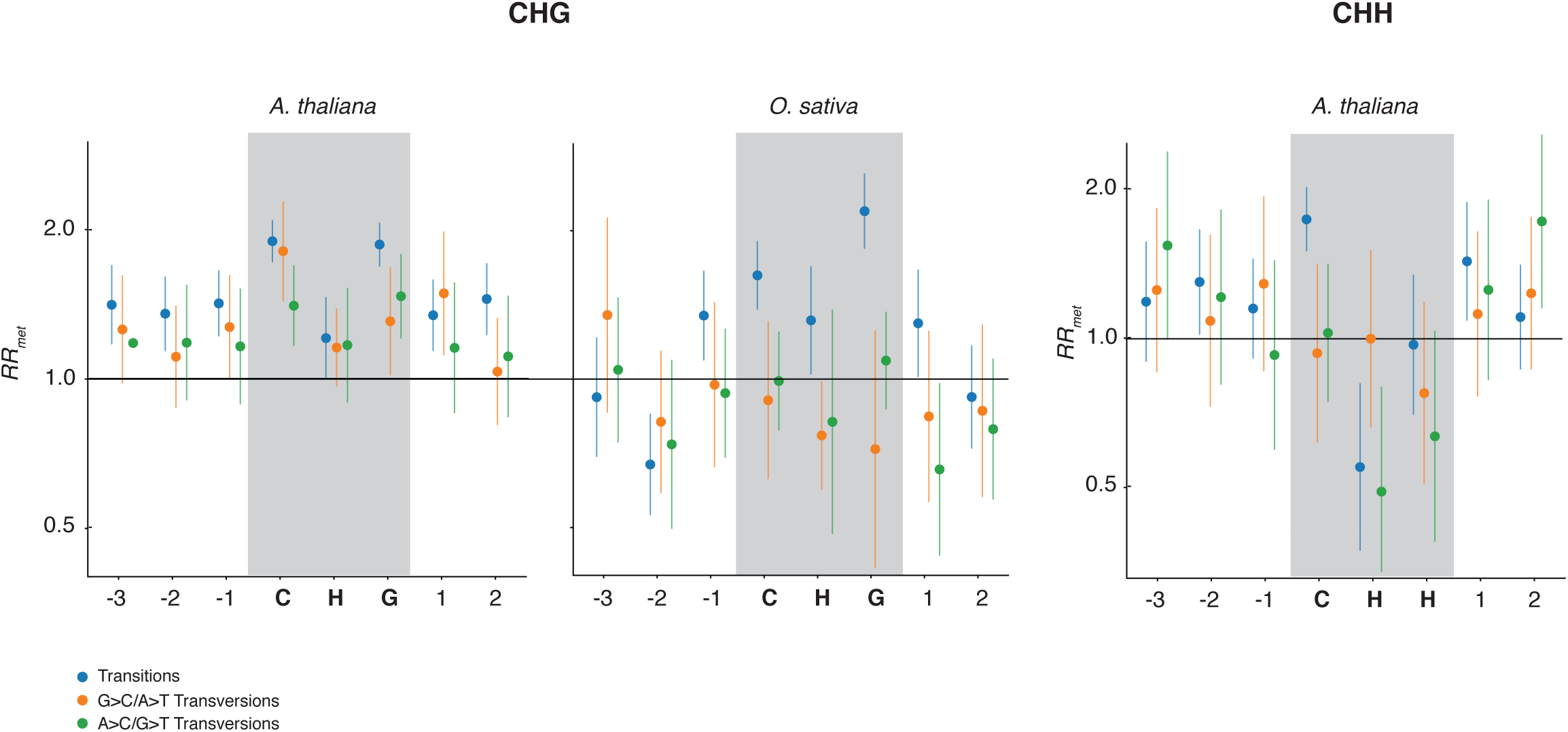
The relative mutational risk of methylation (*RR_met_*) at CpHpG an CpHpH sites and neighbouring nucleotides in *A. thaliana* and rice. *RR_met_* estimates for CpHpH in rice are too noisy to be informative and are therefore not shown. P-values for overall *RR_met_* estimates are: CpHpG (*A. thaliana*): P=3.8*10^−19^; CpHpH (*A. thaliana*): P=1.8*10^−8^; CpHpG (*O. sativa*): P=0.043. Vertical bars are confidence intervals on *RR_met_* computed using the delta method.

**Figure S2.**
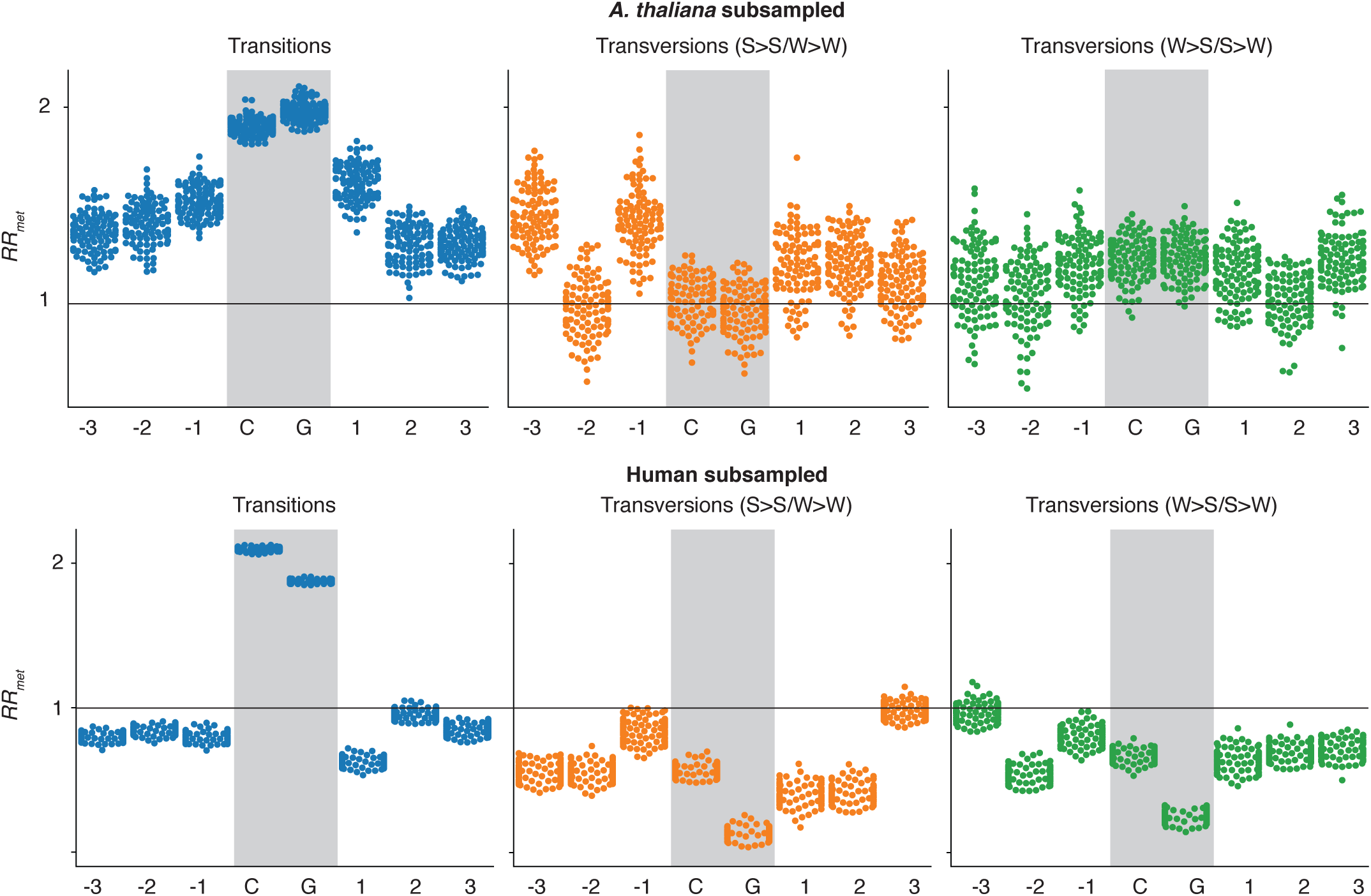
*RR_met_* of subsampled motifs. The number of occurrences for each observed nucleotide motif was counted in both human and *A. thaliana* motif pairs. For each motif, the occurrences in both sets were compared, and the set with the larger number of said motif was randomly subsampled without replacement to contain the same number as the smaller sample. After doing this for all motifs, we have a reduced set of human and *A. thaliana* methylated and unmethylated motifs with identical motif distribution. *RR_met_* was then calculated as described. This process was repeated 100 times, to account for biases in random sampling.

**Figure S3.**
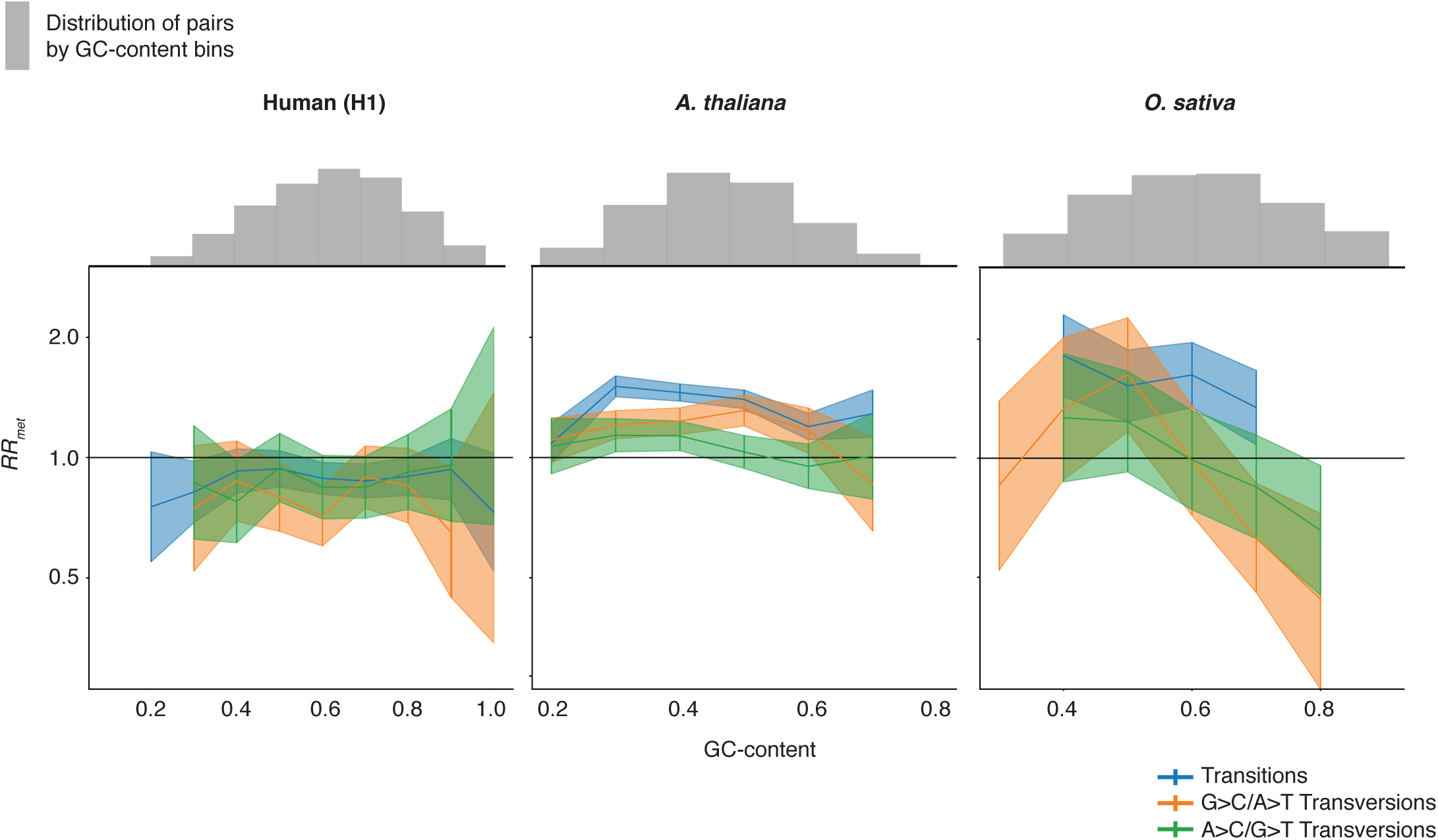
The relative mutational risk of methylation (*RR_met_*) at CpG-neighbouring nucleotides in human, *A. thaliana*, and rice as a function of motif GC content. *RR_met_* was calculated based on GC content bins. *RR_met_* estimates are omitted where error bars were larger than the plotting range. Vertical bars are confidence intervals on *RR_met_* computed using the delta method.

**Figure S4.**
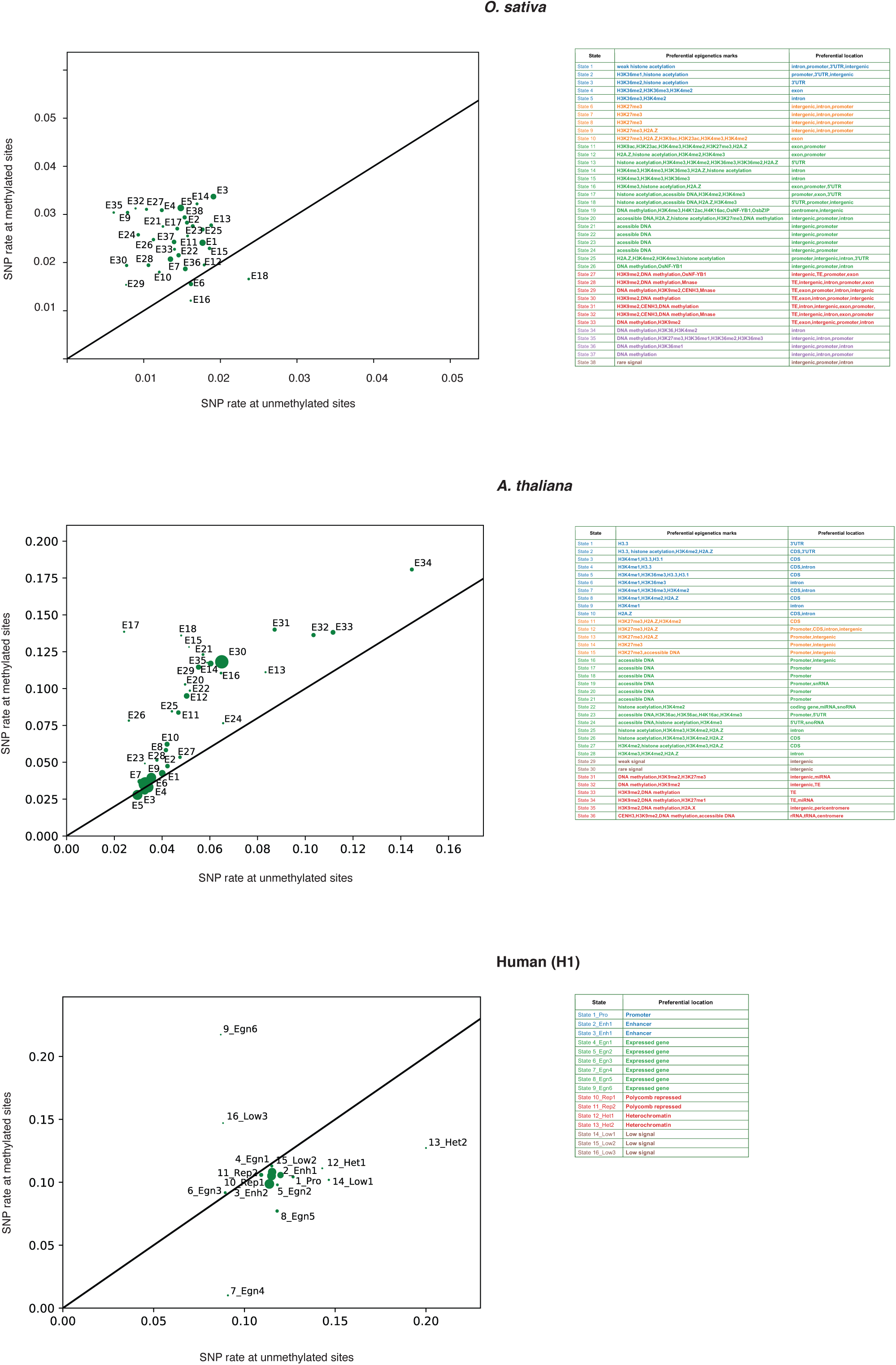
*RR_met_* in *A. thaliana,* human, and rice for different chromatin states, deconstructed into SNP rates at methylated and unmethylated sites across pairs. Individual chromatin states, classified as described in the Methods, are labelled. Text colour corresponds to the colouring of chromatin states in Figure 3.

